# Feature Detection to Segment Cardiomyocyte Nuclei for Investigating Cardiac Contractility

**DOI:** 10.1101/2021.03.03.433810

**Authors:** Tanveer Teranikar, Cameron Villarreal, Nabid Salehin, Jessica Lim, Toluwani Ijaseun, Hung Cao, Cheng–Jen Chuong, Juhyun Lee

## Abstract

*In vivo* quantitative assessment of structural and functional biomarkers is essential for understanding pathophysiology and identifying novel therapies for congenital heart disorders. Cardiac defect analysis through fixed tissue and histology has offered revolutionary insights into the tissue architecture, but section thickness limits the tissue penetration. This study demonstrated the potential of Light Sheet Fluorescence Microscopy (LSFM) for analyzing *in vivo* 4D (3d + time) cardiac contractility. Furthermore, we have described the utility of an improved feature detection framework for localizing cardiomyocyte nuclei in the zebrafish atrium and ventricle. Using the Hessian Difference of Gaussian (HDoG) scale space in conjunction with the watershed algorithm, we were able to quantify a statistically significant increase in cardiomyocyte nuclei count across different developmental stages.

Furthermore, we assessed individual volumes and surface areas for the cardiomyocyte nuclei in the ventricle’s innermost and outermost curvature during cardiac systole and diastole. Using the segmented nuclei volumes from the feature detection, we successfully performed local area ratio analysis to quantify the degree of deformation suffered by the outermost ventricular region compared to the innermost ventricular region. This paper focuses on the merits of our segmentation and demonstrates its efficacy for cell counting and morphology analysis in the presence of anisotropic illumination across the field-of-view (FOV).

## INTRODUCTION

The evolution of optical microscopy has resulted in a high volume of image attributes with differing degrees of complexities^1^. The consequence is a grueling effort to validate the interpretation and reliability of a biomarker based study^2^. Moreover, manual analysis often leads to inconsistencies in the generalization of biomarker studies due to the incredible number of target features compared to sampling size in clinical images^2,3^. In this regard, feature detection methods help refine data dimensionality by discarding dispensable image attributes, thereby producing a more compact feature space^1,2^.

In biomedical research, endogenous fluorescent signals are commonly used to study *in vivo* spatiotemporal organogenesis. However, optical microscopy inherently suffers from anisotropic illumination in the optical image plane^4^. This anisotropy is due to aberrations caused by photon propagation through heterogeneous tissue morphology or significantly varied sample size; hence, optimization of image acquisition parameters is a challenging task^4^. Precise orchestration of any volumetric reconstruction analysis requires high feature detection sensitivity with respect to tissue protrusions and changes in illumination, rotation, local scaling changes, and motion translation^5^. These optical aberrations are prone to induce redundancy in image features^4^, rendering the task of manual feature extraction intrinsically subjective and laborious^6,7^. Hence, feature detection has gained popularity for augmenting performance of downstream classifiers or regression estimators to improve their interpretability and accuracy^8^.

Quantitative zebrafish cardiomyocyte proliferation studies involving embryonic heart volumetric reconstruction have reported strict regulation of regional mechanisms such as increase in cell size and proliferation rate for normal heart maturation and chamber formation^9,10^. Moreover, previous studies have shown hypertrophy in cell size due to older cell differentiation or migration into smaller-sized cardiomyocytes^9^. With recent progress in Light Sheet Fluorescence Microscopy (LSFM), it is possible to perform *in vivo* 4D (3D + time) cardiac cell tracking and perform mechano-transduction studies^11,12,13^. When this method is employed using transgenic zebrafish, optical sections at different tissue depths are acquired to understand the cardiac architecture^11,14,15,16^. However, quantifying *in vivo* cardiac biomechanics remains a challenging task due to improper focus induced by sample movement or sampling artifacts.

Additionally, embryonic zebrafish cardiomyocyte cell counting enables researchers to understand cell proliferation during cardiogenesis^17,18^. However, many studies use manual boundary demarcation due to the limited availability of binary classification methods impervious to heterogenous contrast resolution.^9,19^. Traditionally intensity-based segmentation methods such as the Otsu’s method, adaptive thresholding, iso data thresholding, and entropy-based thresholding have been used for automated cell tracking for their simplicity and speed^20^. However, these methods are infamous for poor performance in the presence of noise and hence incapable of separating objects in close proximity into meaningful biological regions^21^. Consequently, all the thresholding techniques mentioned above suffer in cell detection performance and cannot perceive separate events at different scales, arising from separate biological processes. Additionally, noise due to optical aberrations, short exposure time, cells entering and leaving the field of view and insufficient contrast between adjacent cells may all diminish the technique’s performance^3,21^. Another conventionally favored approach for cell isolation involves the watershed algorithm^3,22,23^. However, the technique is fraught with over-segmentation or false detection of non-existent objects in the presence of noise or complex cell morphology^3,21^. Hence, to suppress noise and reduce the number of false positives with respect to detected cell count after binarization^24^, we used the difference of Gaussian (DoG) scale-space as a pre-processing strategy before contour detection by the Hessian difference of Gaussian (HDoG) operation. We have tried to mimic the human visual perception (HPV) for our feature detection strategy, advertently making it more sensitive to edges than brightness variation^4,25,26^. However, the DoG band-pass operation can induce under-detection of features due to suppression of high frequency noise^27^. Consequently, we performed local curvature evaluation from the Hessian matrix^28^ in conjunction with the HPV based DoG detector, for precise cardiomyocyte nuclei boundary analysis and localizing individual nuclei volumes.

The feature detection framework can be efficiently applied for cell segmentation in developmental biology^3,23^ with post-processing strategies based on mathematical morphology such as opening, closing, top-hat and bottom-hat transform for pruning redundant binary features^4,28^. Contemporary sophistication for dealing with heterogeneity in nuclear volumes involves a predetermined template based approach^29^. It comprises two functions: a marker function to denote object locations and a segmentation function used to delineate the object boundary^3,22,19,30^. Other interest points detectors that effectively deal with scale heterogeneity across the features of interest are scale space methods such as the Hessian blob alogorithm^6,31^.

In this study, we have evaluated the performance of a novel pre-processing framework based on combining scale space and local curvature filters^6,28,31^, in order to separate cardiomyocyte nuclei contours across different scales accurately. We have applied our edge detection method to quantify *in vivo* cardiac contractility by tracking cardiomyocyte nuclei and exhibited the ability of framework for distinguishing individual cardiomyocytes from background noise in the developing zebrafish heart. Moreover, we have validated the accuracy of the feature detection in the presence of real-time imaging artifacts and its stability for ground truth developmental cardiac optical image segmentation for cell tracking.

## RESULTS

### Isolating the coalesced cardiomyocytes from beating zebrafish heart image

Zebrafish cardiogenesis is characterized by augmented cell proliferation and distinct phases of cell differentiation, resulting in the addition of new cardiomyocytes to the heart^17^. Distinguishing the fused cardiomyocyte nuclei volumes by manual hand segmentation or determining the intensity threshold for the region of interest (ROI) suffering from poor edge contrast is an exhausting task due to undiscernible overlapping nuclei boundaries (**Figure 1A**). Furthermore, low sampling rates, endogenous auto-fluorescence, and the beating heart’s dynamic motion convolute the lateral and axial imaging planes making it extremely difficult to obtain pristine images of high optical clarity **(Figure 1B-D)**. To segment the fused cardiomyocyte cluster morphology into disjoint regions while avoiding over segmentation due to non-uniformly illuminated boundaries, we implemented the watershed algorithm after using the difference of Gaussian (DoG) filter to enhance cell boundaries. As a result, we were able to successfully separate closed contour nuclei from each other in a moderately dense cell environment at 48 hpf (**Figure 1E-H, Video S1**). Moreover, we were achieved isotropic segmentation across the lateral and axial imaging planes, respectively, suffered from anisotropic illumination across the field-of-view (FOV) (**Figure 1I-L**).

**Figure 1.**
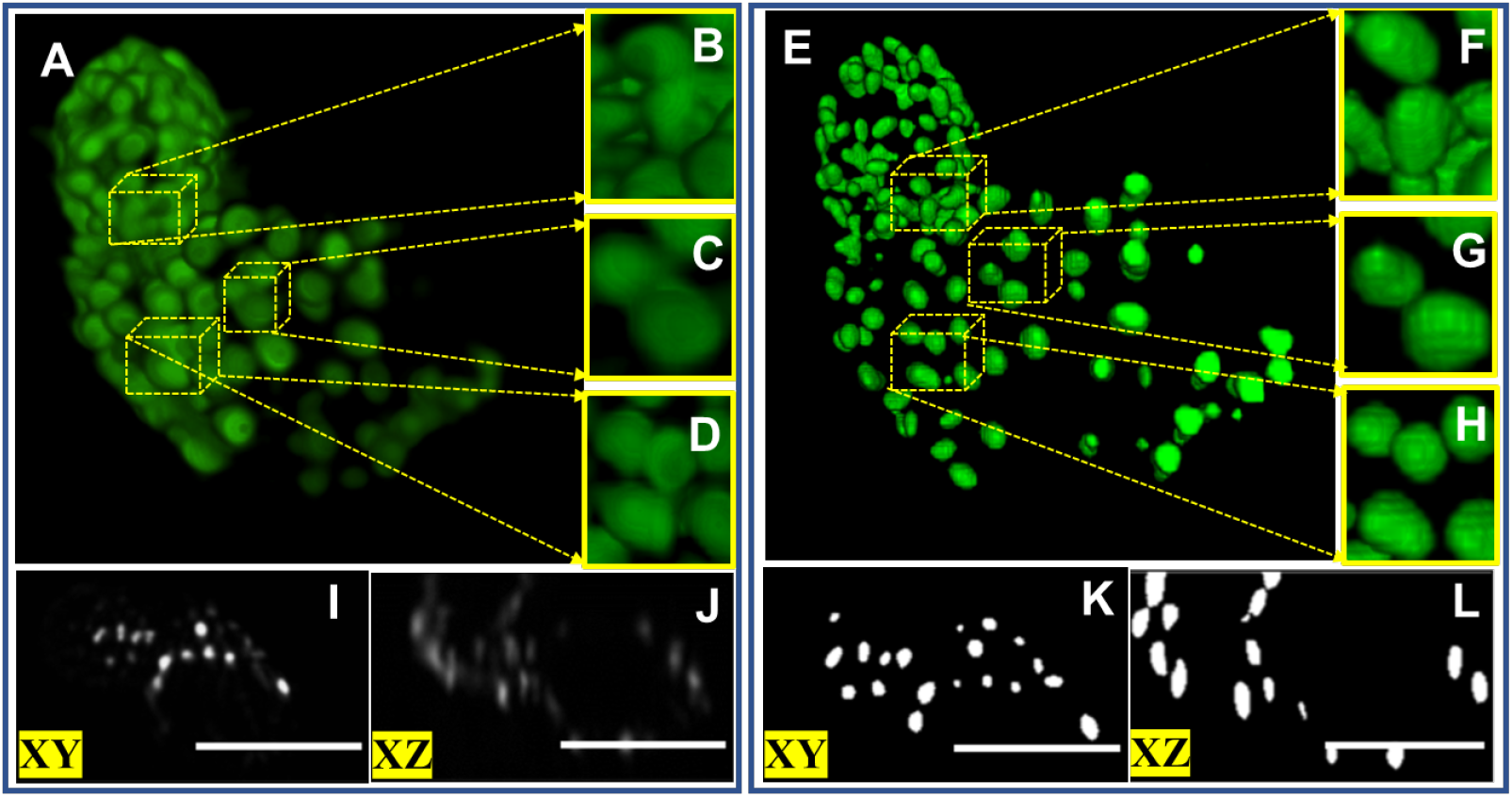
Isolating and segmenting cardiomyocyte nuclei from contracting heart using the DOG (Difference of Gaussian) filter in combination with the watershed algorithm. (A) 48 hpf volumetric reconstruction of light-sheet microscopy acquired image was used to visualize time dependent motion of cardiomyocytes. Raw image was limited in distinguish single cardiomyocyte nuclei from dense environment which visualized at clumped cluster (yellow highlighted boxes). (B-D) Three parts of entire 2 dpf zebrafish heart were zoomed to show that cardiomyocytes were clumped. Thus, tracking or counting cardiomyocytes were hampered. (E) Dynamic volumetric image was processed using the Difference of Gaussian (DoG) blob detector in conjunction with the watershed algorithm to successfully distinguish single cardiomyocytes from adjacent neighbors. Cardiomyocytes in zoomed in areas compared to raw image were distinct and recognize as countable cardiomyocytes (yellow highlighted boxes). (F-H) Three zoomed in parts show that each cardiomyocytes were clearly distinct. (I-J) 2D lateral and axial views show the complex tissue morphology and tissue birefringence in raw images exacerbate image noise, resulting in object overlap. (K-L) Segmented lateral and axial views were visualized as binary data to corroborate curvature delineation of the cardiomyocytes as markers. Scale bar = 50 microns

### Using scale selection for complex cardiomyocyte blob across distinct growth phases

Compared to pinhole-based microscopy techniques, a potential cause of concern for LSFM when imaging zebrafish cardiomyocytes manifests in the form of poor contrast between adjacent cardiomyocytes (**Figure 2A, 2D**)^4^. This poor contrast is due to undesired background fluorescence emitted in regions other than the fluorophore binding sites. Furthermore, intensity attenuation along line-of-sight (LOS) propagation and low Numerical Aperture (NA) can exacerbate the issue of poor signal-to-noise ratio (SNR).

**Figure 2.**
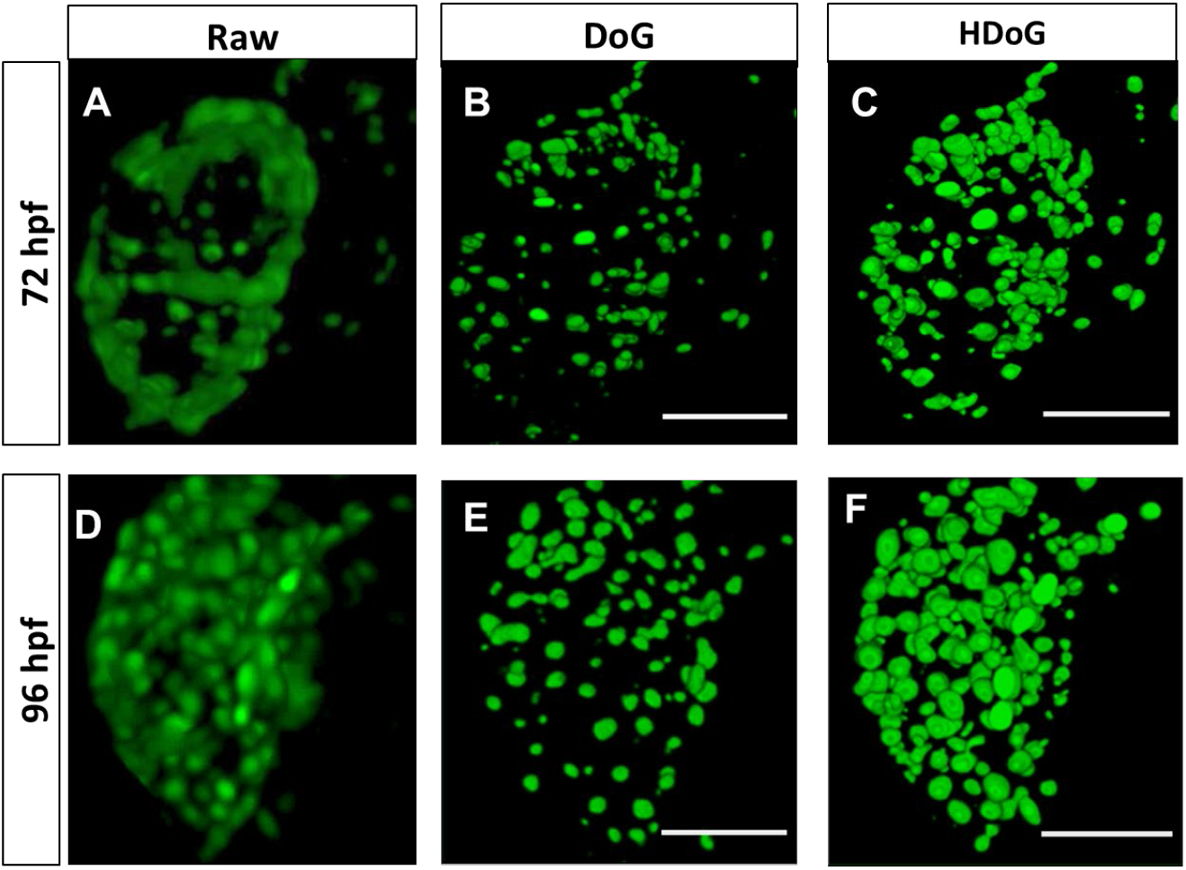
Different blurring scales and imposing boundary conditions on the hessian determinant to separate overlapping cardiomyocyte nuclei and achieve isotropic axial segmentation. (A, D) Raw volumetric reconstruction of 72 hpf and 96 hpf were relatively larger size of zebrafish and more cardiomyocyte numbers compare to 48 hpf. Therefore, tissue scattering circumvents isotropic intensity of fluorescent illumination and causes blurred images. (B, E) 72 hpf and 96 hpf segmented volumes obtained by using the watershed algorithm in combination with the DOG detector limited in under segmentation which caused incorrect tracking or count of cardiomyocytes from dynamic volumetric images due to disappearance during contraction and relaxation. (C, F) Application of Hessian Difference of Gaussian (HDoG) detector to 72 hpf and 96 hpf segmented volumes in conjunction with the watershed algorithm warrants a blob detector that is sensitive to false blob candidates as well as over segmentation. The segmented volumes show improved sensitivity of detector response with respect to local affine transformations experiences by pixel neighborhoods in the image slices such as rotation (sample movement), reflection as well as shear mapping. (scale bar = 50 micron)

Although we distinguished individual atrial and ventricular cardiomyocyte nuclei in a moderately dense cell environment at 72 hpf, we encountered under-segmentation and inaccurate visual due to low pixel intensities produced by the DoG edge detector (**Figure 2B, 2E, and Video S2)**. Therefore, we applied the Hessian Difference of Gaussian detector (HDoG) for accurate cell count and morphology measurements of cardiomyocyte nuclei in a dense environment. We used the hessian scale space representation to localize saddle points^6^. Saddle points are points in a function that are neither an intensity maximum nor minimum, that can represent connecting nuclei edges. This approach improves detection sensitivity in the presence of multiple intensity peaks for a single biomarker and for exact biomarker boundary detection for feature analysis (**Figure 2C, F, and Video S3**).

### Segmentation accuracy evaluation

We first visualized and quantified the zebrafish contractile motion in 4-D image at different scales, for each distinct developmental phase: 48 hpf (**Figure 3A – 3D**), 72 hpf (**Figure 3E – 3H**) and 120 hpf (**Figure 3I – 3L**). We used a segmentation ratio to test the accuracy of the segmentation compared to the raw images of the cardiomyocytes, which also served to evaluate segmentation accuracy. In this case, the segmentation ratio is the number of segmented cardiomyocytes divided by the number of cardiomyocytes in the raw images. If the numerical value = 1, the segmentation is identical to the raw images. If the numerical value > 1, there is over-segmentation in the segmented images. If the numerical value < 1, there is under segmentation in the segmented images. Our analysis found that the ideal segmentation score was repeated across all developmental stages using the Hessian characteristic scale space with DoG detected features as blobs (**Figure S1**).

**Figure 3.**
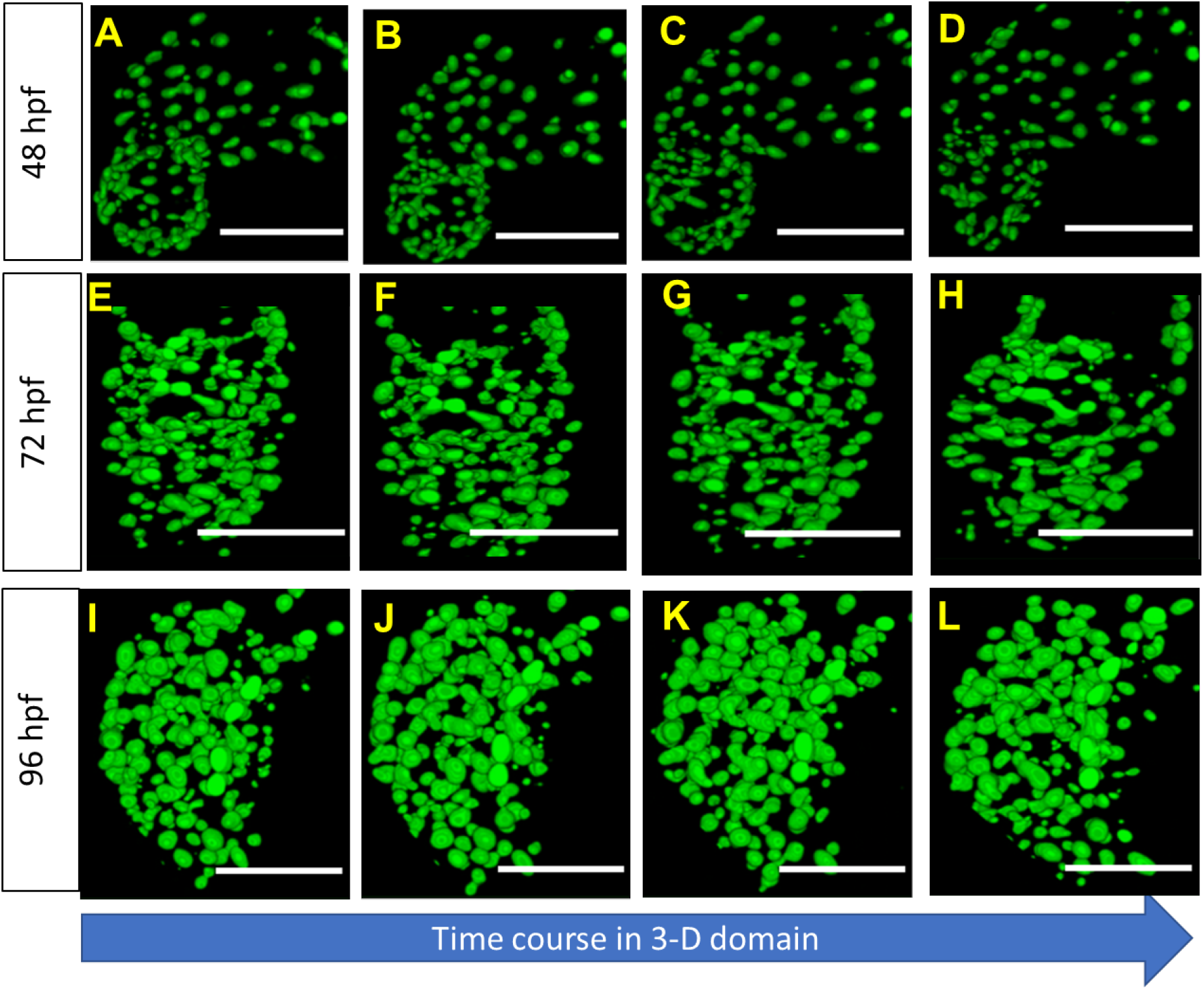
Varying tissue morphology across different developmental stages affects cardiomyocyte separation. (A-D) cardiomyocytes were distributed throughout dynamic zebrafish heart under looping condition at 48 hpf (E-H) At 72 hpf, individual cardiomyocyte nuclei was visualized. Despite of anisotropic Gaussian luminance across different developmental stages and motion of cardiomyocytes, HDoG detector was able to split cardiomyocyte nuclei accurately. (I-L) Even with rapid frame rates required for sampling the contractility in conjunction with, our image processing was able to detect and isolate the myocardial cardiomyocyte nuclei, and subsequently use the segmented labels to assess contractility of the ventricular myocardial cardiomyocyte nuclei. (scale bar = 50 micron)

### Quantification of local contracility via tracking cardiomyocyte nuclei

After we processed the image, we performed contractility analysis by tracking cardiomyocyte nuclei across 48 to 120 hpf to validate this novel segmentation approach (**Figure 4A-H**). We normalized the time course of stretch values for the innermost and outermost curvatures for each developmental stage with respect to the start of end-systolic stage of ventricle (**Figure S2**). We investigated the stretch level change of developing zebrafish heart. In addition to stretch, we investigated area ratio comparison between innermost curvature and outermost curvature areas. Furthermore, the area ratio is a description of an area’s total deformation based on the stretch ratio of two dimensions of the area. We analyzed the time course of area ratio among three cardiomyocytes as markers. We found area ratio of the outermost curvature area, where the opposite side of the atrioventricular canal receiving the pumped blood directly from the atrium, has a higher area ratio than the innermost curvature area of the ventricle. However, both areas increase the area ratio consistently (**Figure 4I-J**).

**Figure 4.**
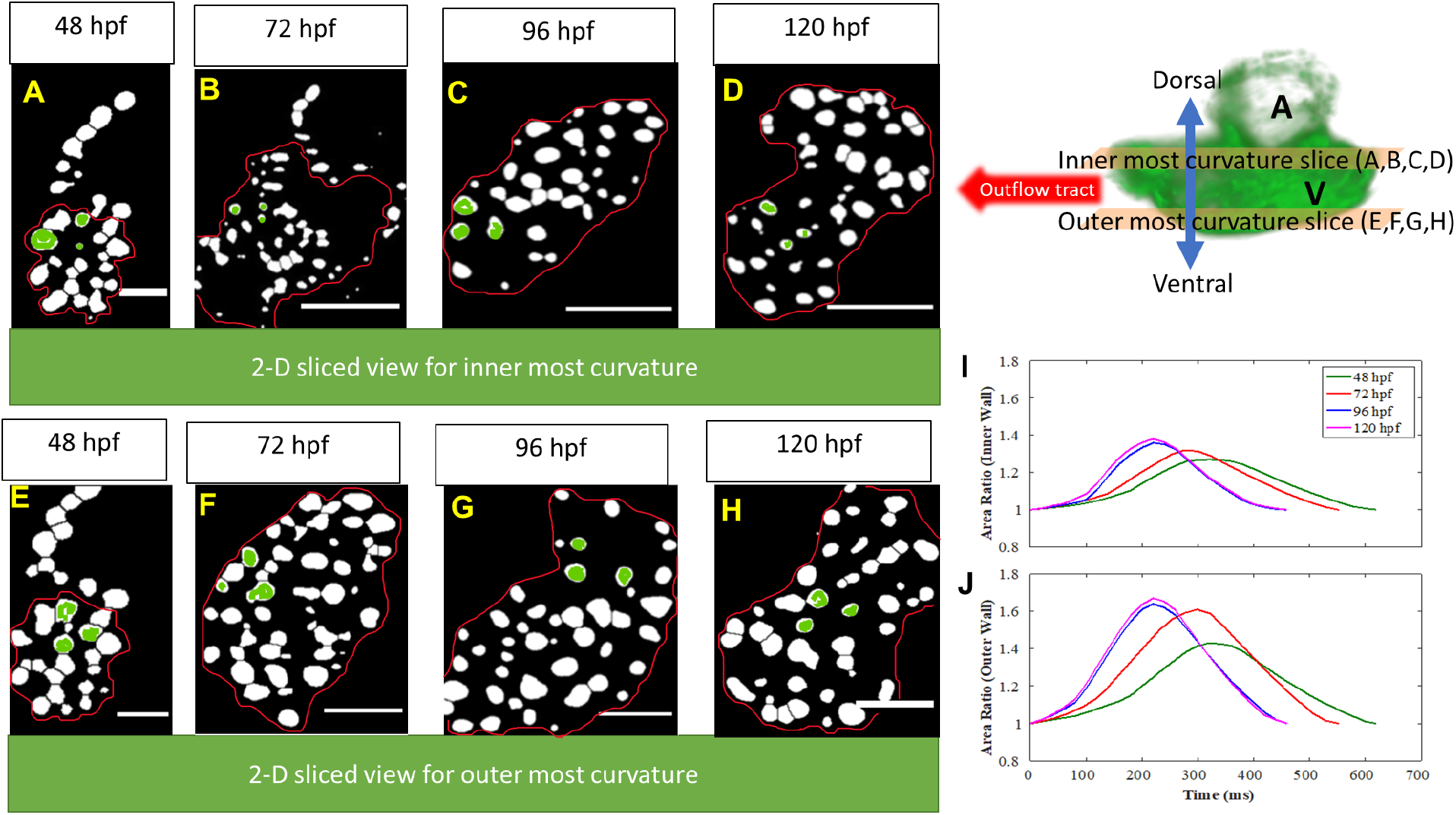
Selected markers utilized area ratio analysis. (A-D) 2-D sliced views of 48 hpg, 72 hpf, 96 hpf, and 120 hpf of zebrafish dorsal area of ventricle featured inner most curvature, atrioventricular canal. **(E-H)** 2-D sliced views of 48 hpf, 72 hpf, 96 hpf, and 120 hpf of zebrafish ventral part of zebrafish heart featuring outer most curvature of ventricle. **(I)** Area ratio for innermost curvature by tracking three cardiomyocytes highlighted in (A-D) elucidated that increasing trend of contractility by developing zebrafish heart. **(J)** The area ratio of outermost curvature calculated by tracking three cardiomyocytes in (E-H) demonstrated outermost curvature has higher contractility compare to innermost curvature. scale bar for (A) and (E) = 30 microns, scale bar for (B – D), (F – H) = 50 microns.

### Quantification of zebrafish cardiomyocyte nuclei development

The average values for number of nuclei in a developing zebrafish heart were 159±13.1, 222.86 ± 17.50, and 260 ± 13.58, and 284.2±10.844, for 48, 72, 96, and 120 hpf respectively (**Figure 5A, Table S1**) (n = 15. We detected statistically significant differences across all four development stages for nuclei volumes in the zebrafish ventricle (**Table S1**).

**Figure 5:**
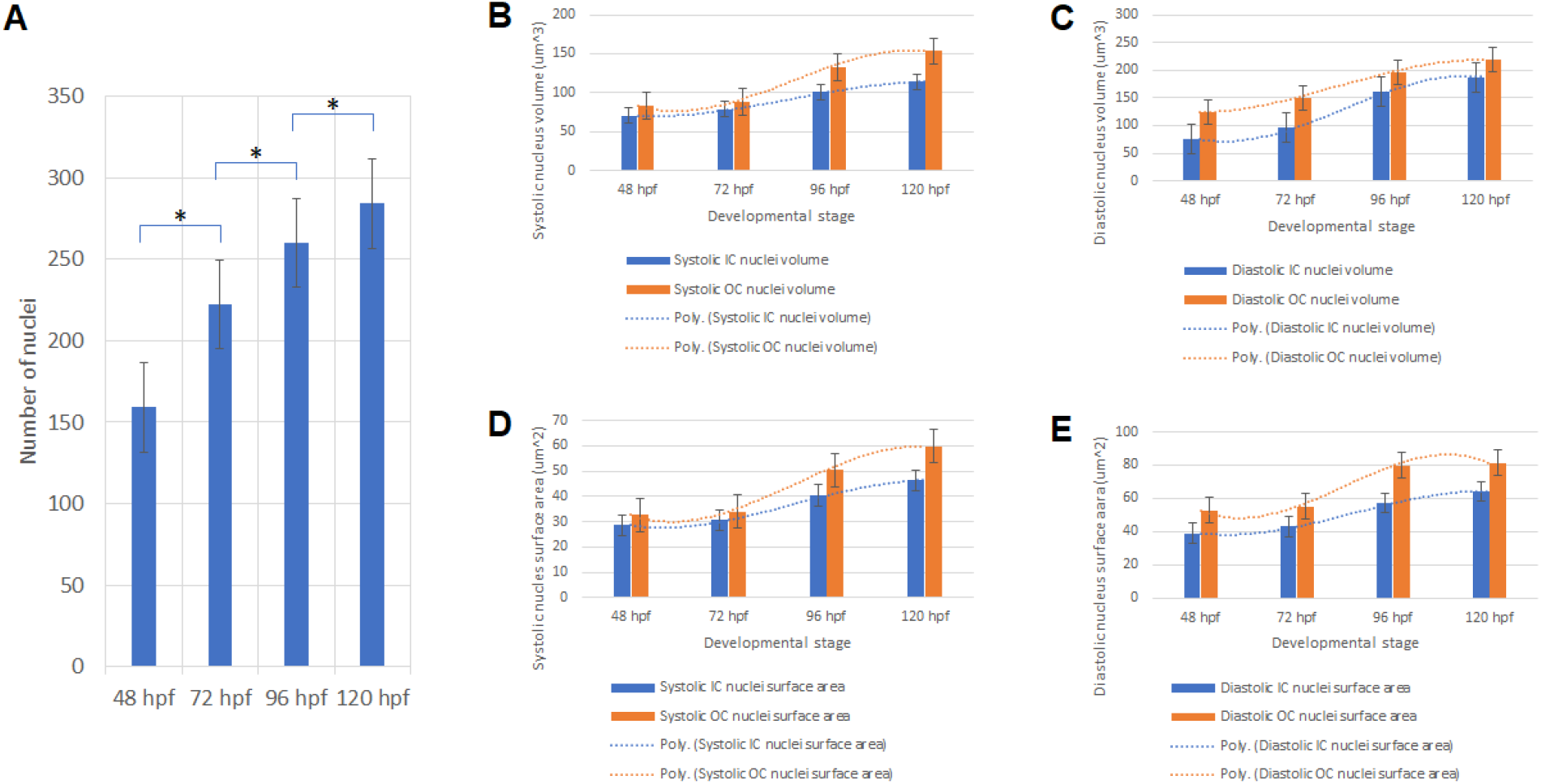
Zebrafish cardiomyocyte nuclei analysis: (A) We observed an increase in the number of ventricular cardiomyocyte nuclei for successive developmental stages. Asterisk denotes statistically significant difference with respect to previous time point. p≤ 0.05 (B) Multinomial trend observed for the systolic nucleus volume in the inner and outer ventricular curvature. (C) Multinomial trends observed for the diastolic nucleus volume in the inner and outer ventricular curvature. (D) Multinomial trend observed for the systolic nucleus surface area in the inner and outer ventricular curvature. (E) Multinomial trend observed for the diastolic nucleus volume in the inner and outer ventricular curvature. n=15

We further investigated systolic and diastolic single nuclei volumes and surface areas for the inner and outer ventricular curvature region (**Figure 5B – 5E**). The multinomial trends indicate a statistically significant rise in systolic and diastolic area measurements for the inner and outer curvature regions respectively, during the 72 – 96 hpf developmental stage only (**Table 1**-**2**). After comparing the inner curvature systolic and diastolic nucleus morphology to the outermost curvature features, we observed a statistically significant difference for the diastolic area measurements across all developmental stages (**Table S2**) but observed significant rise in the systolic area measurements only after 96 hpf (**Table S2**).

**Table 1.** Nucleus volume change between systole and diastole phases.

|  | Nucleus Systolic volume |  | Nucleus Diastolic volume |  |
| --- | --- | --- | --- | --- |
|  | Inner curvature | Outer curvature | Inner curvature | Outer curvature |
| **48 hpf vs 72 hpf** | p = 0.4 | p = 0.6 | p = 0.1 | p = 0.1 |
| **72 hpf vs 96 hpf** | P = 0.05 * | P = 0.001* | P = 7.79 E<sup>-05</sup>* | P = 0.04* |
| **96 hpf vs 120 hpf** | P = 0.2 | P = 0.12 | P = 0.06 | P = 0.3 |

**Table 2.** Nucleus surface area change between systole and diastole phases.

|  | Nucleus Systolic surface area |  | Nucleus Diastolic surface area |  |
| --- | --- | --- | --- | --- |
|  | Inner curvature | Outer curvature | Inner curvature | Outer curvature |
| **48 hpf vs 72 hpf** | P = 0.6 | P = 0.6 | P = 0.2 | P = 0.7 |
| **72 hpf vs 96 hpf** | P = 0.01 * | P = 0.001 * | P = 0.01 * | P = 0.001 * |
| **96 hpf vs 120 hpf** | P = 0.1 | P = 0.1 | P = 0.2 | P = 0.8 |

As a result, we were able to successfully localize the fluorescent biomarkers to quantify the number of cells in the zebrafish ventricle and the ventricular inflow and outflow regions. Furthermore, we were assessed the area characteristics of individual segmented nuclei volumes during the systole and diastole, for the beforementioned ventricular regions.

## DISCUSSION

Cardiomyocytes are ellipsoid shaped units that occupy significant myocardial architectural volume and are fundamentally responsible for myocardial contractility^33,34^. Precise contractile evaluation of the myocardial contractility requires isolation of the individual cardiomyocytes from the surrounding tissue^34^. Current myocardial deformation methods involving cardiomyocyte volume isolation by heart extraction of rat, mouse, or guinea pig models suffer from the limited investigation into tissue due to histological section thickness^33^. Consequently, statistical significance is hard to prove due to the limited sampling size and lifetime of viable cells^33^. However, recent advancements in non-invasive cardiac optical sectioning microscopy^11,13,18,35^ and transgenic zebrafish generation have enabled accurate *in vivo* 4D cardiac contractility quantification by cardiomyocyte nuclei tracking analysis^18,35^.

In this study, we evaluated a novel approach based on well-established methods for robust biomarker edge detection. We successfully quantified the zebrafish ventricular myocardial contractility using the segmented nuclei at distinct developmental phases. Moreover, our improved HDoG edge detection framework provides direct access to surface area and volume measurements of fluorescently labeled cardiac nuclei to report accurate cell morphology characteristics. Although we induced transparency in the zebrafish embryos, tissue birefringence causes heterogeneous refractive index^4^. This briefcasing could severely affect the incoming light sheet’s tissue penetration ability, resulting in anisotropic edge illumination across the field-of-view^4^. Taking this into consideration, we sought to design a feature detector that would provide robust performance for cardiomyocyte nuclei detection accuracy with respect to 1) dynamic volume contraction, 2) single local maxima generation for a true biomarker by Hessian, and 3) localization of segment to the nuclei center by DoG. Other framework design constraints involved consistency in feature detection with respect to illumination, scale (nuclei area heterogeneity) and rotation invariance. Previously conducted studies have reported the use of Laplacian of Gaussian (LoG) and Difference of Gaussian (DoG) feature detectors for efficiently separating individual cell proliferation from a dense environment^6,31^. We have used the computationally inexpensive DoG as a Gaussian filter based tunable bandpass operation for enhancing edge visibility for our study. The DoG blob detector involves identifying the optimal degree of specimen blurring that provides maximum filter response for isolation of local maxima of different sizes. Moreover, the DoG filter also functions as a greyscale edge enhancement algorithm, that is useful for images experiencing refocus. From our analysis, the algorithm is capable of handling the varying nuclei orientation and heterogenous nuclei sizes (**Figure 1**) but still suffered under reporting of edges in high cell density environment (**Figure 2B, 2E**). The filter response for locating blob center is dependent on matching the blurring scale and scale (width) of the interest blob. Intuitively, the scale space representation image can be understood as a 2D slice in which all objects with dimensions matching the blurring scale (standard deviation of gaussian blur) are wiped out^6^. Hence, the scale space is a hierarchal family of 2D images from a lower to higher blur degree, with each scale space representation image having a distinct number of edges. A single pixel counts as theV lowest limit, all the image pixels together constitute the upper limit of feature scale^6^. HDoG scale space filtering thus involves qualitative description of the image features and subsets of the original image are obtained by convolving the raw image with gaussian blur masks of different orders^36^.

Our binary thresholding used for separating touching nuclei will prove beneficial not only for counting the varying number of cardiomyocytes but also for all types of cells to understand the rate of cell proliferation. Additionally, analyzing the distinct cardiomyocyte motion (**Video S4**) allowed us to quantify local deformation to understand the mechanotransduction of biological signaling during development. For example, we demonstrated that the outermost curvature has higher area ratio than the innermost curvature (**Figure 4I-J**). Indeed, these morphological parameters are evidence of cardiac maturation, and further studies can utilize the segmentation method developed in this study to elaborate on these findings.

In summary, we have presented a blob detection approach for segmenting a dense nuclear network, with individual nuclei exhibiting distinct morphological characteristics. DoG blob detection followed by boundary segmentation using the watershed algorithm is insufficient for precise contour description of the touching cardiac myocyte nuclei edges. Undesirable computational inconsistencies are induced due to inability of filter response to localize multiple intensity peaks in the same ROI. Hence, to avoid nuclei undercount and inaccurate volume measurement, we combined the DoG and Hessian scale space representation prior to intensity thresholding. Our blob detection framework provides robust performance for images corrupted with noise or insufficient depth detail. With our proposed framework’s help, biomechanical quantification of a developing zebrafish heart was performed, along with accurate counting of touching nuclei to understand the cell proliferation rate.

## METHODS

### Light sheet microscope (LSFM) implementation

Our home built tunable LSFM consists of a single side illumination pathway and a water dipping lens (insert lens description) detection setup^4^. In the illumination pathway, an optical train consisting of a cylindrical lens in conjunction with a 4x objective lens (4X Plan Apochromat Plan N, Olympus, Tokyo, Japan) is used to produce a collimated cylindrical light-sheet with Gaussian intensity distribution. We adjusted the mechanical slit space to cover the circumferential ventricular volume across 48 - 120 hours post fertilization. As the sample is scanned along the axial direction, the optical detection train consisting of a sCMOS camera (ORCA flash 4.0, Hamamatsu, Japan), coupled with a water dipping lens, is used to acquire real time 4D (3D + time) cardiac atrial and ventricular volumes as described previously^30,31^.

### Preparation of zebrafish for assessing cardiac function

The animal experiments were performed in agreement with the UT Arlington Institutional Animal Care and Use Committee (IACUC) protocol (#A17.014). The transgenic zebrafish line used in this particular study is the *Tg*(*cmlc2*:*nucGFP*), with the cardiac myocyte nuclei labeled with GFP (Green Fluorescent Protein)^19^. The zebrafish embryos were maintained at 28.5°C in system water at the UT Arlington Aquatic Animal Core Facility. 0.0025% 1-phenyl 2-thoiurea was added to the embryo medium starting at 20-24 hpf ^4,32^ to suppress pigmentation. Prior to imaging, the embryos were anesthetized in 0.05% tricaine and embedded in 0.5% low-melt agarose, within a refractive index matching (RIM) fluorinated ethylene propylene (FEP) tube (refractive index of water = 1.33, refractive index of agarose and FEP tube = 1.34). The mounting scheme involved suspension of FEP tube into a 3d printed enclosure with RIM cover slip windows, for non - invasive tissue penetration.

### Image processing framework

Supplementary documents describes detail information of image processing framework step-by-step. In addition, image processing code is enclosed (**Code S1**).

### Cell counting and area measurements

After converting raw optical images to binary images, we performed to count cardiomyocyte nuclei and their area analysis by using the 3D object counter plugin^33^ in ImageJ. The plugin can be accessed by: ImageJ – Analyze – 3D Object Counter. After cropping the ROI (ventricle in this case), we used the plugin to quantify number of object voxels (volume), surface voxels of individual nuclei volumes and the number of 3D nuclei objects in the ventricular stack. The plugin can also be used to retrieve the centroid geometric coordinates of object volumes. The user is required configure 2 important parameters namely, (a) intensity threshold to separate background and foreground pixel populations and (b) size threshold to exclude smaller objects from the analysis. The plugin allows user to configure object counting based on the presence or absence of touching edges.

### Cardiac myocyte nuclei tracking

We utilized the segmented, processed, and time synchronized images to reconstruct three-dimensional volumes through time for a cardiac cycle to perform this tracking. We then passed these images through a custom MATLAB (Mathworks) code to perform for key steps(**Code S2**).

### Contractility Analysis

We selected and tracked three cardiomyocyte nuclei for both the innermost and outermost curvature. After tracking the location of three cardiomyocytes through each time instance, we utilized the following method to determine the deformation gradient with normalized one cardiac cycle as 0.5 second starting from ventricular end-systolic stage. We determined the stretch ratio at each time instance into principal stretch values reported as λ_1_ and λ_2_, or the longitudinal and circumferential principal direction followed by previous methods^41,42^. These principal stretch vectors correspond to the first and second principal strain directions. When viewed on Mohr’s circle, they correspond to the maximum and minimum normal strain values where the shear strain is resolved to zero (**Figure S4**). These values are represented in the Cartesian coordinate system as the x and y direction or in polar coordinates as the zero-degree rotation and 90-degree rotation. We established the area ratio by multiplying the two principal stretch values. Area ratio provides a description of the total in plane deformation from the initial undeformed state which was selected as the start of filling.

### Statistics

For statistical analysis, we performed ad hoc pairwise comparisons for three morphological parameters to characterize the maturity of the heart^10^. We analyzed the number of visible nuclei, the total volume, and total surface area. We estimated each of these parameters using built in functions in ImageJ (NIH, Bethesda, MD) with n=15. Additionally, we cleaned the data in excel utilizing Chauvenet’s criterion to determine which values were outliers and should be removed. After removing outliers and cleaning the data sets in excel to reduce the chance of error due to our sampling technique, we compared the data with one-way ANOVA. If we detected a statistically significant difference for any comparison, we performed Tukey’s test for multiple comparison of means. This test inherently compensates for multiple comparisons, which allowed us to use an alpha value of 0.5. All values herein are reported as mean plus or minus standard deviation.

## ACKNOWLEDGEMENTS

The authors would like to express gratitude to Dr. Caroline Burns and Geoffrey Burns from Boston Children’s Hospital for providing *tg(cmlc:nucGFP)* for imaging and analysis. This study was supported by grants from AHA 18CDA34110150 (J.L.) and NSF 1936519 (J.L.).

