## Supplementary document for "Feature Detection to Segment Cardiomyocyte Nuclei for Investigating Cardiac Contractility"

### Supplementary Text

This document is used to describe the mathematical equations for the preprocessing workflow described in the Results and Methods section. References have been provided at the end of the document.

**Supplementary information list**

**Figure S1:** Quantifying segmentation accuracy
**Figure S2:** Time course of the principal stretch values for developing zebrafish
**Figure S3:** Global ventricular cardiomyocyte nuclei morphology
**Figure S4:** Tracking cardiomyocyte nuclei for assessing cardiac contractility across different developmental stages
**Table S1:** Statistical significance of tested parameters
**Table S2:** Statistical significance of ventricular cardiomyocyte nucleus at systole and diastole for the inner and outermost curvature region. Asterisk denotes significant change from previous value
**Video S1:** 2dpf zebrafish heart (DoG→watershed)
**Video S2:** 3dpf zebrafish heart (DoG→watershed)
**Video S3:** 3dpf zebrafish heart (Hessian→DoG→watershed)
**Video S4:** 4 dpf zebrafish heart (Hessian→DoG→watershed)
**Video S5:** Resolved planar motion through one cardiac cycle
**Code S1:** Image processing framework
**Code S2:** Cardiomyocyte motion tracking code

**Image processing Framework**

**Figure S1**

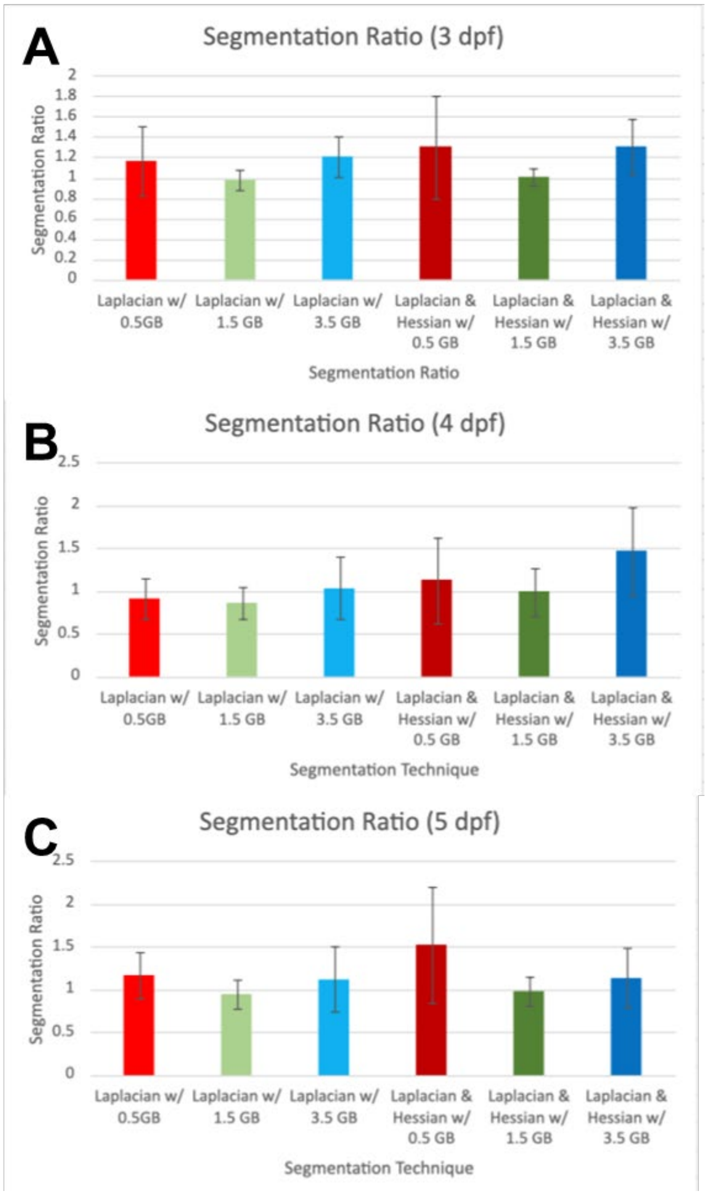


**Figure S1.** Quantifying segmentation accuracy. We tested the precision of the proposed framework to assess false positives or negatives that may be introduced as markers. We constructed scale spaces for the Difference of Gaussian Filter and the hessian + DoG respectively , by blurring volumes at different degrees of blur ( 0.5 – 3.5 Gaussian kernel). By identifying the characteristic scale space representation (1.5 Gaussian blur), we observed a perfect segmentation result with respect to visually inspected nuclei count across different developmental stages. (GB = Gaussian blurring degree)

**Figure S2.**

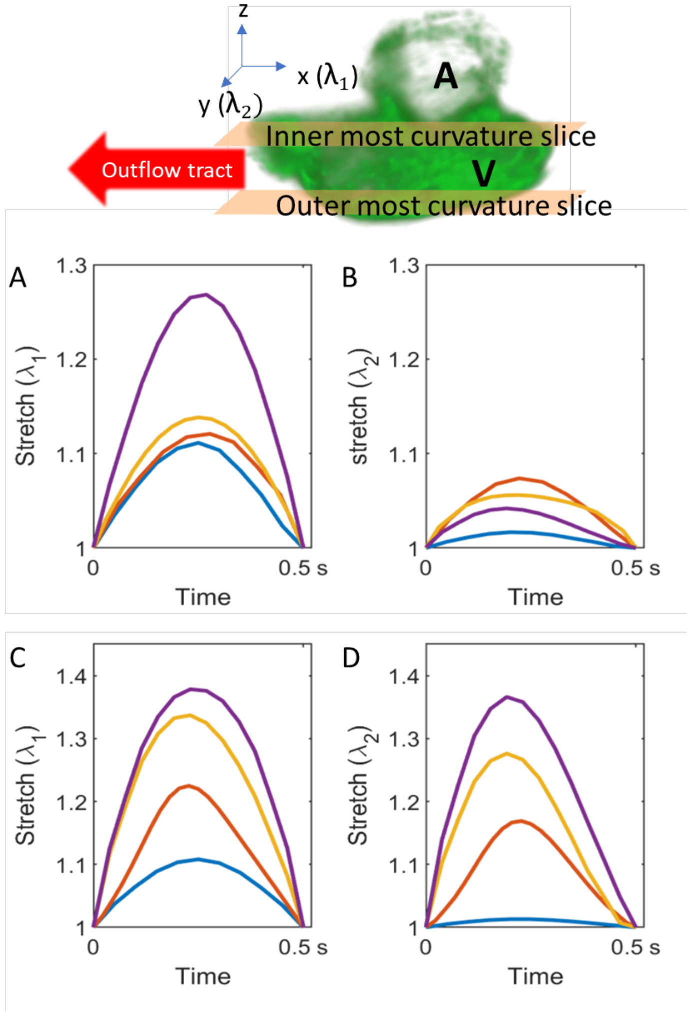


**Figure S2.** The graphs represent the time course of strain in principal directions one and two for the ventricle's outermost and innermost curvature. Where epsilon 1 and epsilon 2 are relative to the changing orientation of the triangular plane utilized in analysis. Relative to each configuration λ1 lies in the longitudinal direction and λ2 lies along the circumferential direction. The start of ventricular filling is the reference point for analysis. At this stage, the ventricle has the lowest volume, and the change in stretch through the cardiac cycle is more evident. The graphs correspond to the following values: (A) Innermost curvature stretch in longitudinal direction, (B) innermost curvature stretch in circumferential direction, (C) outermost curvature in longitudinal direction, and (D) Outermost curvature in circumferential direction.

**Figure S3.**

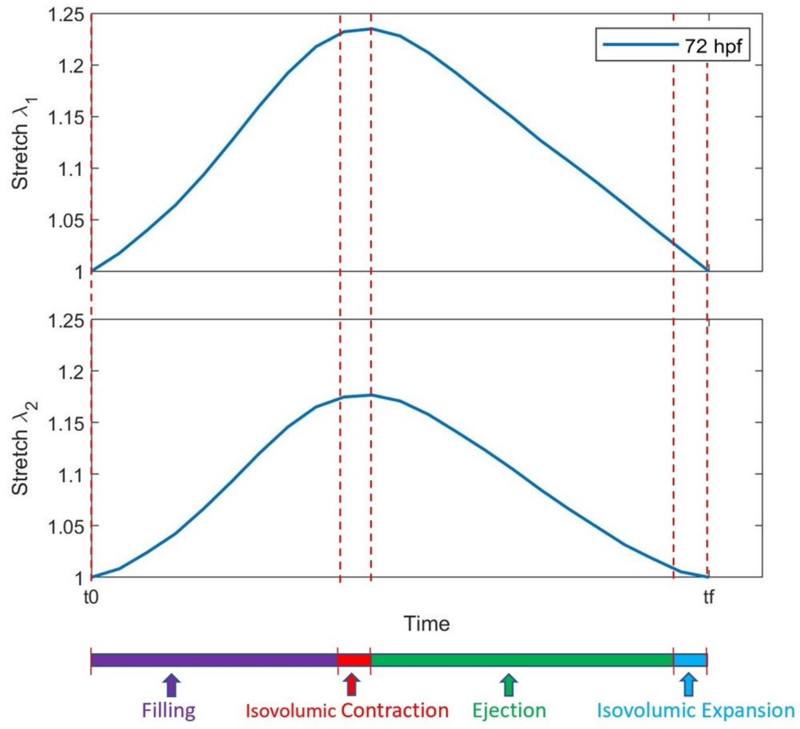


**Figure S3.** Time course of the principal stretch values for 3dpf zebrafish taken from the outermost curvature with a reference state of the lowest ventricular volume when ventricular filling begins. This figure displays key steps in the cardiac cycle superimposed on the time course of principal stretch values lambda 1 and lambda 2. Critical points of interest include ventricular: filling, isovolumic contraction, ejection, and isovolumic expansion.

**Table S1.** Statistical significance of tested parameters

| Comparison | P-value | Parameter |
|---|---|---|
| 48-72 | 0.001 | Number of Cardiomyocyte Nuclei |
| 48-96 | 0.001 | |
| 48-120 | 0.001 | |
| 72-96 | 0.001 | |
| 72-120 | 0.001 | |
| 96-120 | 0.001 | |
| **Comparison** | **P-Value** | **Parameter** |
| 48-72 | 0.001 | Surface Area |
| 48-96 | 0.001 | |
| 48-120 | 0.001 | |
| 72-96 | 0.0621 | |
| 72-120 | 0.001 | |
| 96-120 | 0.001 | |
| **Comparison** | **P-Value** | **Parameter** |
| 48-72 | 0.001 | Volume |
| 48-96 | 0.001 | |
| 48-120 | 0.001 | |
| 72-96 | 0.0177 | |
| 72-120 | 0.001 | |
| 96-120 | 0.001 | |

**Table S2**: Statistical significance of ventricular cardiomyocyte nucleus at systole and diastole for the inner and outermost curvature region. Asterisk denotes significant change from previous value.

| | **Nucleus systolic volume** | **Nucleus diastolic volume** |
|---|---|---|
| | Inner curvature vs Outer curvature | Inner curvature vs Outer curvature |
| 48 hpf | P = 0.2 | P = 0.0003 |
| 72 hpf | P = 0.4 | P = 0.01* |
| 96 hpf | P = 0.002* | P = 0.03* |
| 120 hpf | P = 0.008* | P = 0.04* |

| | **Nucleus systolic surface area** | **Nucleus diastolic surface area** |
|---|---|---|
| | Inner curvature vs Outer curvature | Inner curvature vs Outer curvature |
| 48 hpf | P = 0.3 | P = 0.01 |
| 72 hpf | P = 0.3 | P = 0.03* |
| 96 hpf | P = 0.03* | P = 0.001* |
| 120 hpf | P = 0.02* | P = 0.01* |

**Code S1.** Image processing framework (Please see detail description in the bottom page)

**Code S2.** Cardiomyocyte motion tracking code

This Matlab code performed following 4 steps.

1. The code compiles images into easily searchable 4D matrices.
2. The code resolves the 4D matrices of segmented images into centers of mass based on high pixel concentration areas for each time step.
3. The user selects three markers to represent our plane for stretch calculations.
4. The code searches through the 4D stack of centers of mass to determine the closest center of mass in the next time step and stores these points in a matrix of position values.

Each stored triplet value is the x, y, and z position of a particular nucleus at a particular time. This format is easily searchable and allows for a multitude of calculations. This code assumes that there can be no erratic motion of the nucleus with a high enough sampling frequency. The location at each time step depends on the prior location. Imaging with a high sampling frequency will supply data that meets this assumption requirement. Other works have utilized similar works, including Meijerling *et al* [1]. Drawbacks of this method include the requirement for user interaction. To verify that the cell tracking occurs appropriately, the user must analyze each vector to ensure the vector does not violate the small motion assumption. This process can become time-consuming and increases the chance of human error. Subsequent work can expand and refine this cell tracking method to include other parameters, including a probability net for machine learning applications and size and orientation to decrease ambiguity and reduce the user input requirement

### Image processing Framework

*Haze Removal using the Dark Channel Prior (DCP) method*

Introduction of haze by the ambient medium or scattering due to particulate matter, degrades the performance of computer vision tasks[2,3]. A haze free image can be retrieved by using the image degradation model based on the Dark Channel Prior (DCP) algorithm,

$$I(x)=J(x).t(x)+A(1-t(x))^{2,3} - (1),$$

Where I(x) is the degraded image, J(x) is the original irradiance captured by the CMOS camera, t(x) represents the scene depth and A is the scattering introduced by the ambient light. Using the dehazing algorithm, we estimated the intensity transmission map t(x) using the imreducehaze() matlab function[3].

$$t(x) = e^{-\beta d(x)2} - (2)$$

where $\beta$ represents the scattering coefficient and d represents the scene depth. We used the estimated intensity transmission map as a preprocessing step before performing the DoG operation. By estimating the contrast attenuation with respect to distance, we were able to emphasize edges.

*Intensity maxima localization at nuclei centers using the Difference of Gaussian (DoG) filter*

The DoG filter can be effectively used to enhance edge visualization for images suffering from poor contrast. In this study, the greyscale bandpass operation is performed by subtracting a blurred version of the transmission estimate from a lesser blurred version of itself,

$$t(x)^* g1(x) - t(x)^* g2(x) = t(x)^* (g1(x) - g2(x)) - (3),$$

where g1(x) and g2(x) are the gaussian kernels having different standard deviations. Using the DoG filter, we were able to localize blobs to nuclei centers by isolating spatial frequencies correlating to the gaussian illumination maxima.

*Precise contour delineation using the hessian scale space representation and watershed algorithm*

The hessian scale space representation can be described by the convolution:

$$D(x,y,t) = [t(x)* (g1(x) – g2(x))] * G(x,y,t)$$ [4,5] – (4),

Where D(x,y,t) represents the family of images , derived from the original image. t represents the degree of blurring. Hence, equation (4) can be described as the convolution of the DoG blob maxima image with the hessian blob detector gaussian blur kernel G(x,y) at different degrees of blur ($t > 0$). The blurring scale selection was based on the ratio $t +1 = r * t$ [4], where r is a constant.

The workflow involved for the hessian blob involves[4,5],

1) Computing the absolute magnitude of the intensity gradient image obtained by convolving the DoG bandpass image with the derivative of gaussian filter.
2) Computing the double derivative of the absolute magnitude image calculated in the previous step
3) Imposing boundary conditions on the hessian determinant value [$\det D(x,y,t) < 0$][5] at every pixel , for indicating saddle points.

The image arithmetic operation (OR – operation) results in the union of the DoG localized intensity maxima and contour information from the Hessian blob, aiding the successful splitting of nuclei.

*Preprocessing strategies*

Images corrupted by noise or tissue scatter, were filtered by using a gaussian kernel with an appropriate standard deviation followed by the background subtract operation in ImageJ.
